# Multi-Task Learning based 3-Dimensional Striatal Segmentation of MRI – a Multi-modal Objective Assessment

**DOI:** 10.1101/2020.07.13.200576

**Authors:** Mario Serrano-Sosa, Jared X. Van Snellenberg, Jiayan Meng, Karl Spuhler, Jodi J. Weinstein, Anissa Abi-Dargham, Mark Slifstein, Chuan Huang

## Abstract

The human striatum is a collection of subcortical nuclei that serves as a critical node in an information processing network. Recent studies have established a clear topographical and functional organization of projections to and from striatal regions that involves detailed and complex subdivisions of the striatum. Manual segmentation of these functional subdivisions is labor-intensive and time-consuming, yet automated methods have also been a challenging computational problem. Recently, deep learning has emerged as a powerful tool for various tasks, including semantic segmentation. Multi-Task Learning (MTL) is a machine learning technique that allows latent representations between related tasks to be shared during model prediction of multiple tasks with better accuracy. We utilized MTL to segment subregions of the striatum consisting of pre-commissural putamen (prePU), pre-commissural caudate (preCA), post-commissural putamen (postPU), post-commissural caudate (postCA), and ventral striatum (VST). Dice similarity coefficients (DSC) demonstrate excellent spatial agreement between manual and MTL-generated segmentations (≥ 0.72 across all striatal subregions). Further quantitative task-based analysis was also conducted. Binding potential values, BP_ND_, of [11C]raclopride PET, and ROI time-series and whole-brain connectivity using fMRI images were compared between results generated from manual segmentations and MTL-generated segmentations. BP_ND_ values from MTL-generated segmentations were shown to correlate well with manual segmentations with R_2_ ≥ 0.91 in all caudate and putamen subregions, and R_2_=0.69 in VST. Mean Pearson correlation coefficients of the fMRI data between MTL-generated and manual segmentations were also high in time-series (≥0.86) and whole-brain connectivity (≥0.89) across all subregions. We conclude that the MTL approach is a fast, robust and reliable method for 3D striatal subregion segmentation with results comparable to manually segmented ROIs.

## 1. Introduction

The human striatum is a collection of subcortical nuclei that serves as a critical node in an information processing network proceeding from cortex, through the striatum and other basal ganglia structures (e.g. globus pallidus and subthalamic nucleus), to the thalamus and back to cortex. Striatal pathology or dysfunction is thought to play a critical role in neurological and neuropsychiatric diseases such as schizophrenia, Parkinson’s, and Huntington’s disease_1–4_. Functional and molecular imaging studies of patients with these disorders frequently require identifying the portion of an image volume that contains the striatum and then segmenting the striatum into its anatomical or functional subregions _5,6_. Automated segmentation of the striatum has been a challenging computational problem, and some researchers continue to rely on manual delineations that are labor-intensive and time-consuming. A plethora of automatic and semi-automatic methods have been proposed to segment brain regions including the striatum. Widely available atlas-based approaches, such as Freesurfer, can robustly identify the whole striatum and its major structures (the caudate and putamen), but have been less reliable in segmenting the striatal subregions, particularly in patient populations_7_.

More recently, deep learning has emerged as a powerful tool for various tasks, including semantic segmentation. Deep learning has the capacity to learn hierarchical features in a data-driven manner through its numerous convolutional neural network (CNN) layers that represent different levels of abstraction. Automated deep learning techniques for semantic segmentation of brain regions from Magnetic Resonance Imaging (MRI) has been recently used to segment subcortical regions, including the striatum and its major structures, the caudate and putamen_8,9_; however, performance was mainly evaluated using Dice Similarity Score (DSC) with no task-based analysis to determine the accuracy of their segmentations for research use. This latter deep learning method_9_ for striatal segmentation also used two separate networks, a Global CNN to localize the striatum and a Local CNN to further confirm if the chosen voxels were striatal. Although this method provided decent results, more robust deep learning methods exist that do not require multi-network segmentation. Specifically, segmentation using the U-Net architecture has been shown to capture complex feature context from the contracting path and to enable precise localization from the symmetric expanding path_10_. More importantly, no deep learning technique for subregion segmentation has been reported yet.

Anatomical tracing studies in nonhuman primates have established a clear topographical and functional organization of projections to and from striatal regions that involves more detailed and complex subdivisions of the striatum than gross anatomical divisions would suggest_11_, such that there may be clear benefits to neuroimaging studies if this topography can be captured and utilized in data analysis. Imaging researchers, particularly in positron emission tomography (PET), have translated this topographical organization into operationally defined substructures that can be identified and manually traced on T1-weighted or multi-modal MR images_12,13_. These approaches require some visual identification and manual outlining of substructures by analysts with expert knowledge and are time consuming, requiring a day or more to complete a single striatum. As a result, functional Magnetic Resonance Imaging (fMRI) studies very rarely, if ever, use manual segmentation, with most studies employing a set of several “seed” regions of interest that are implemented as small spheres drawn in a standardized coordinate space in the striatum_14–18_. However, this approach fails to take any account individual differences in striatal topography and may fail to optimally capture meaningful subdivisions in striatal structures.

Multi-Task Learning (MTL) is a technique used in deep learning that allows representations between related tasks to be shared that can generalize the model to predict the original task, and many more, with better accuracy_19_. Also called joint learning, learning to learn, and learning with auxiliary tasks, this method has been known to optimize multiple tasks and reduce overfitting_19,20_. In this study we sought to utilize MTL to segment subregions of the striatum consisting of pre-commissural putamen (prePU), pre-commissural caudate (preCA), post-commissural putamen (postPU), post-commissural caudate (postCA), and ventral striatum (VST) using a convolutional neural network (CNN). More specifically, we employ a 3D U-Net architecture with single input and multiple outputs to segment the striatal regions of interest (ROIs) and background. Here, we assess these striatal segmentations with DSC with respect to the gold standard manually drawn ROIs. More importantly, we use task-based analysis applied to the MTL segmentations to re-analyze previously published PET binding potential data, as well as resting state functional connectivity (RSFC) fMRI data, obtained from manually drawn ROIs_21_ and compare these results to those obtained with our new methodology.

## 2. Methods

### 2.1 Datasets

#### 2.1.1 Training/Validation Data

Sixty-eight data sets from both previously published_22,23_ and ongoing studies of patients with schizophrenia and matched controls were employed. Striatal and extrastriatal ROIs were manually drawn on individual T1-weighted scans using a previously validated method_24,25_, which is a time-consuming procedure. The 3D T1-weighted image was used as input for the MTL network and six output tasks were simultaneously trained; where five of the six were ROI masks and the last was the masked background. 68 total datasets were split into 64 training and 4 validation sets (See Table 1).

**Table 1.** Details of the T1-weight MRI data used in this study.

|  | Training set 1 | Training set 2 | testing set |
| --- | --- | --- | --- |
| Sample size | 46* | 22* | 19 |
| Age (SD) | 32.5(10.2) | 31.0 (12.1) | 33.2(9.7) |
| Sex (M/F) | 21/25 | 15/7 | 13/6 |
| Scanner | Philips Intera 1.5T | GE MR 750 3T | GE MR 750 3T |
| Original publication | Horga et al., 2016;<br>Van Snellenberg et al.,<br>2016<sup>22,23</sup> | Ongoing,<br>unpublished | Weinstein et al.,<br>2018<sup>21</sup> |
| Sequence | SPGR SENSE | 3D BRAVO | 3D BRAVO |
| TR (ms) | 25 | 7.82 | 7.82 |
| TE (ms) | 5.67 | 3.09 | 3.09 |
| Parallel imaging factor | 1.5 | 2 | 2 |
| Voxel size (mm) | 1 | 0.8 | 0.8 |
| Flip angle | 30 | 12 | 12 |

\* 1 and 3 subjects out of training set 1 and 2, respectively, were used as validation

**Table 1:** Mean Dice Similarity Coefficient (DSC) with standard deviations for all sub-region ROIs.

|  | prePU | preCA | postPU | postCA | VST |
| --- | --- | --- | --- | --- | --- |
| **DSC** | $0.8123 \pm$ | $0.8716 \pm$ | $0.8234 \pm$ | $0.7506 \pm$ | $0.7221 \pm$ |
|  | 0.0795 | 0.0364 | 0.0348 | 0.0485 | 0.0643 |

#### 2.1.2 PET Test Data set

We used data from a previously published study examining dopamine D2 receptor internalization in 10 patients with schizophrenia and 9 matched healthy controls_21_, which were independent from the training datasets. PET data were acquired on a Siemens mCT scanner. T1-weighted anatomical images for ROI delineation were acquired for each subject on a GE750 3T scanner. The PET data were corrected for motion and coregistered to the subjects’ MRI data using the normalized maximization of mutual information algorithm in SPM software (Wellcome Trust, Cambridge, UK). ROIs were manually delineated on the MR images as in Mawlawi et al._12_ and applied to the coregistered PET. 60 min of dynamic emission data were acquired, following a bolus injection of 349 ± 109 MBq of [11C]raclopride. Data were reconstructed by filtered back projection (FBP) with computed tomography (CT) used for attenuation correction.

#### 2.1.3 fMRI Test Dataset

A subset of 16 participants (7 patients and 9 controls) from the PET test dataset described above also underwent 30 minutes of RSFC scanning and made up the fMRI test dataset. The fMRI scanning was conducted on the same GE MR 750 3T scanner. Multiband blood oxygen level dependent (BOLD) MR sequences were acquired using a multiband acceleration factor of 6, no in-plane acceleration or parallel imaging, with 66 slices, 192 mm field-of-view, 2 mm isotropic voxel size, 60° flip angle, 850 ms TR, and 25 ms TE. Each participant was scanned in four runs with alternating AP and PA phase encode directions, with each run consisting of 532 volumes. B_0_ field maps were acquired along with gradient echo images in both phase encode directions in the same geometry as the BOLD sequences (B_0_ maps were preferentially used for distortion correction, however in cases where distortion correction failed gradient echo images were used instead).

### 2.2 Network architecture

As shown in Figure 1, this MTL network consisted of a 3D U-Net architecture with an encoding path (contracting, left side) and decoding path (expanding, right side). Images were initially cropped to ensure that the striatum was used as input. The encoding path conforms to the typical architecture of a convolutional neural network (CNN), consisting of the repeated application of three 3 × 3 × 3 convolution layers, each followed by an exponential linear activation (elu). Each block ended with a 2 × 2 × 2 maxpooling layer with stride 2 for downsampling. In addition, at each downsampling step, the number of feature channels was doubled in the encoder path; this increment was initialized at 16 feature maps in the first convolutional block and increased to a maximum of 128. The decoding path consisted of similar convolutional blocks with three 3 × 3 × 3 convolutional layers ending with elu but contain 3 × 3 × 3 convolutional transpose with stride of 2 for upsampling instead of the previously used maxpooling. In this decoding path, the feature channel was halved and skip connection with the corresponding linked feature map from the encoding path was utilized. The final step was a 1 × 1 × 1 convolution that maps the output classification.

**Figure 1:**
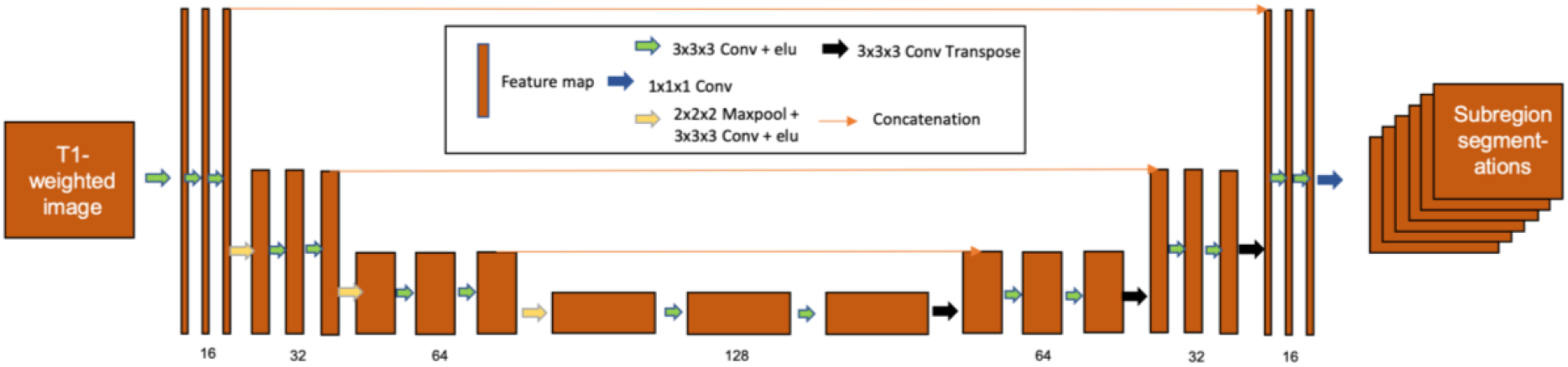
Schematic of the U-Net architecture used for this Multi-Task Learning based segmentation.

### 2.3 Model Training

This model was trained to minimize the sparse softmax cross entropy between output and manually drawn ROIs; where the function first uses softmax to transform the outputs into a probability distribution and computes the loss function:

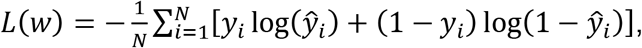

where *W* refers to the model parameters, e.g. weights of the neural network, y_*i*_ is the true label, and 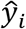 is the predicted label.

The training set and a validation set, together consisting of 68 datasets, was used to optimize the hyperparameters of the model. The model was trained using the Adam optimizer and non-decaying learning rate of 1*10_−5_; network parameters were initialized using the Glorot method_26_. The model was trained on a computer with an i7-8700K 6-core processor, 32 GB memory, and GTX1080 Ti graphics cards running Ubuntu 18.04, Python 2.7.15, and Tensorflow 1.11.0.

### 2.4 Segmentation Evaluation

DSC was utilized to initially evaluate the performance of the MTL network output:

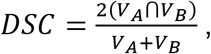

where A is the automatic segmentation from the MTL network and B is the hand drawn ROI.

### 2.5 Task based Assessment

After generating striatal subregion segmentations from MTL network on the test data sets, task-based analyses were conducted on PET and fMRI imaging to determine the reliability of MTL-generated compared to manual segmentations.

#### 2.5.1 Quantitative PET Analysis

In accordance with the original publication_21_ of the PET test set, the primary outcome measure was BP_ND_, the binding potential with respect to the non-displaceable compartment_27_. BP_ND_ was assessed in each ROI using the simplified reference tissue model_28_ (SRTM) with cerebellum as a reference tissue. Here, we compared baseline BP_ND_ values in all 19 subjects from Weinstein et al._21_ to BP_ND_ as assessed using MTL-generated ROIs applied to the subjects’ T1-weighted images and the original coregistered PET data. The kinetic modeling procedure was the same as in the original publication. Linear Regression was performed with BP_ND_ obtained from manual and MTL-generated ROIs. R_2_ and percent differences are reported for manual and MTL-generated ROIs. Intraclass correlation coefficients (ICC(3,2): 2-way mixed model, two-rater, consistency) were also calculated for all subregions with BP_ND_ from MTL-generated segmentations relative to BP_ND_ from manual segmentations.

#### 2.5.2 fMRI Connectivity Analysis

RSFC data was initially preprocessed using the Human Connectome Project (HCP) minimal preprocessing pipeline _29_, and the first 7 volumes of each run were discarded to reach steady-state. Data from one patient participant could not be appropriately distortion-corrected, leaving substantial spatial distortions in the data relative to other participants, and so data from this participant was excluded. Both sets of CNNs were coregistered to RSFC data using SPM 12 spatial coregistration of individual subject T1w images used for manual tracings to the individual-subject space T1w images output by the HCP pipeline. Raw timeseries were then extracted from both hand-drawn and MTL-generated ROIs following preprocessing, and the correlation between the hand-drawn and CNN timeseries data was calculated for each fMRI run. These correlations were then transformed using Fisher’s r-to-Z transformation, averaged across runs, and converted back to r values for presentation.

Next, whole-brain data in MNI space was spatially smoothed using a 4 mm full-width-half-maximum (FWHM) kernel in SPM 12 and underwent standard RSFC processing along with each ROI timeseries in a custom Matlab pipeline: 1) linear detrending and de-meaning of each timeseries, 2) regression of average gray matter, white matter, and CSF signals, 3) regression of motion parameters, their squares, their first derivatives, and their squared first derivatives (“Friston-24” motion parameter regression_30_), 4) identification of “bad” volumes for censoring using recently-developed methods for multiband RSFC data that we have described in detail elsewhere_31_, 5) linear interpolation of signal over bad volumes, 6) bandpass filtering from 0.09 – 0.008 Hz, 7) discarding of the first and last 30 volumes of each timeseries to eliminate distortion due to digital filtering, and finally 8) volume censoring. Timeseries correlations between each voxel were calculated for each ROI and averaged across runs to obtain whole-brain RSFC correlations. Finally, the correlation across voxels in RSFC was calculated between hand-drawn and MTL-generated ROIs.

## 3. Results

The CNN was successfully trained to segment all ROIs. Relative to manually based segmentations that take 1.5 days per subject, our MTL-generated segmentation takes 30 seconds to generate five 3D masks. Figure 2 shows an example of both manual and MTL-generated segmentations; visually, MTL-generated segmentations are similar to manually segmented ROIs. As shown in Figure 2, MTL-generated segmentation seems to smooth VST where manual segmentation may have sharp edges.

**Figure 2:**
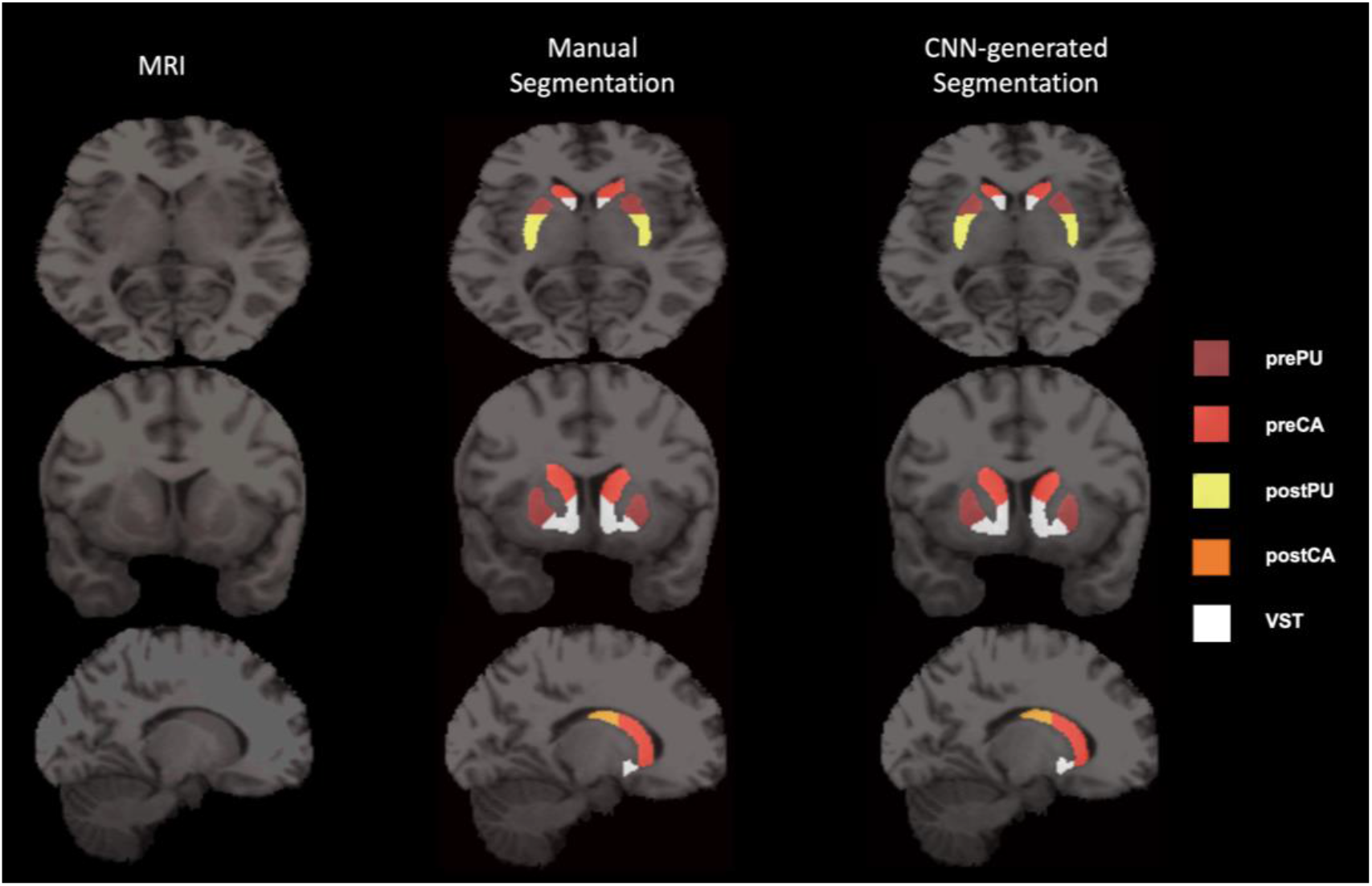
Transverse (top), coronal (middle) and sagittal (bottom) views of T1-weighted MR Image (left), overlaid with ROIs by manual segmentation (middle) and MTL-generated segmentation (right). Transverse slices of manual segmentation show left postPU to have irregular shape compared to a smoother left postPU in MTL-generated segmentation. Coronal and sagittal slices of the MRI and manual segmentation also show VST to have irregular shape compared to a smoothed and more circular VST in MTL-generated.

We first tested our MTL-generated segmentations by comparing to manually drawn ROIs using DSC on a completely independent testing set. Table 1 shows mean DSC across the independent testing set with 0.8123 ± 0.0795, 0.8716 ±; 0.0364, 0.8234 ± 0.0348, 0.7506 ± 0.0485, 0.7221 ± 0.0643, for prePU, preCA, postPU, postCA and VST, respectively.

PET quantitative analysis showed R_2_ between BP_ND_ values using manual and MTL-generated ROIs was ≥ 0.91 in all subdivisions of the caudate and putamen, and 0.69 in VST (Figure 3). Table 2 shows regression coefficients for this analysis. The slopes were all close to 1, and the intercepts were relatively small. Figure 4 shows the percent differences of BP_ND_ between the manual and MTL-generated ROIs. Although CNN segmentation of postCA had regression coefficient of 1 and a high R_2_ of 0.91, Figure 4 shows that it had a downward bias. ICC for all subregions was also ≥ 0.835 as shown in table 2.

**Figure 3:**
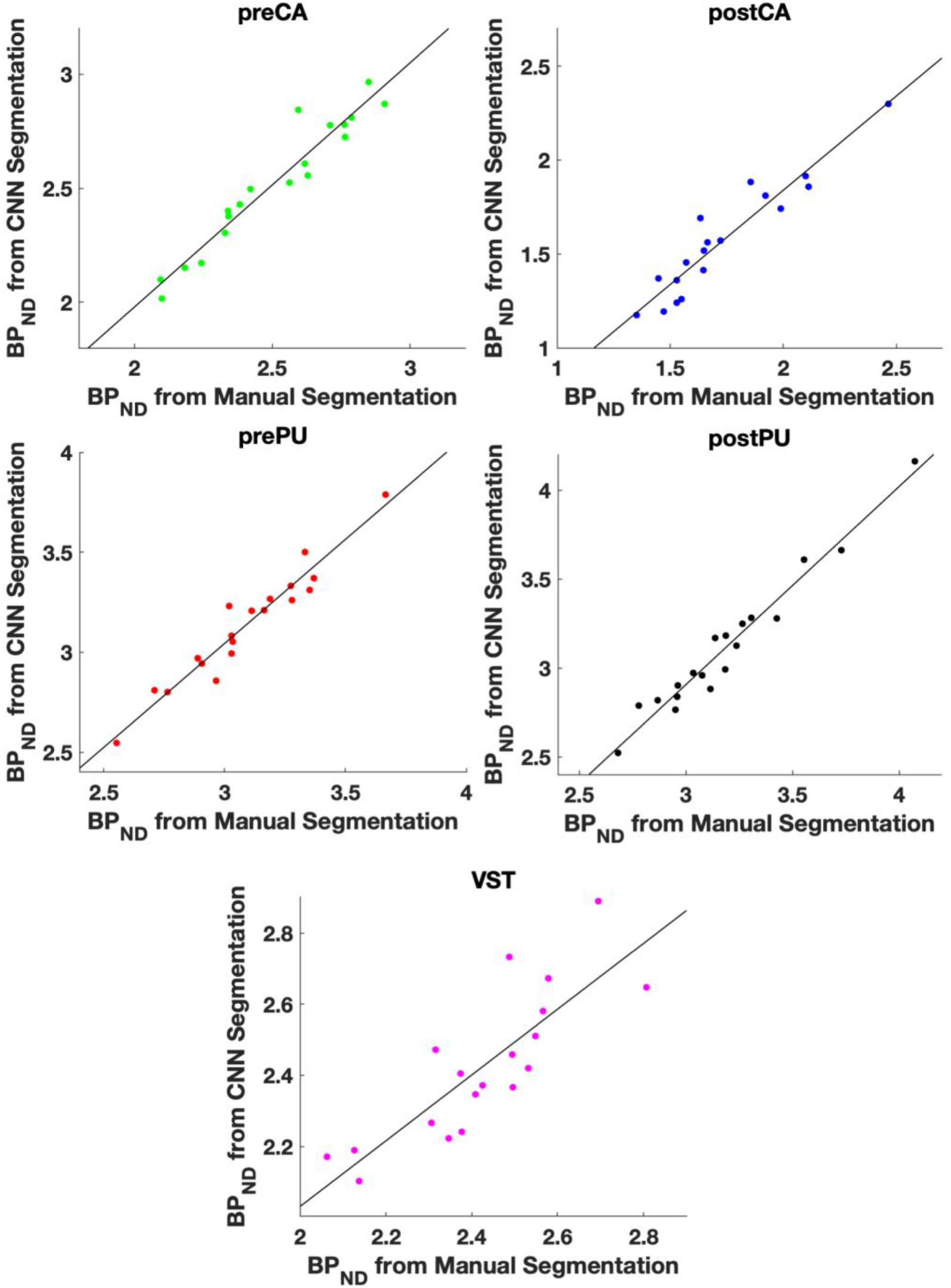
Scatter plots between BP_ND_ of the 5 subregions calculated using manual ROIs and MTL-generated ROIs across 19 independent test subjects. Regression line for all subregions is overlaid scatter plots, with regression coefficients found in Table 2.

**Figure 4:**
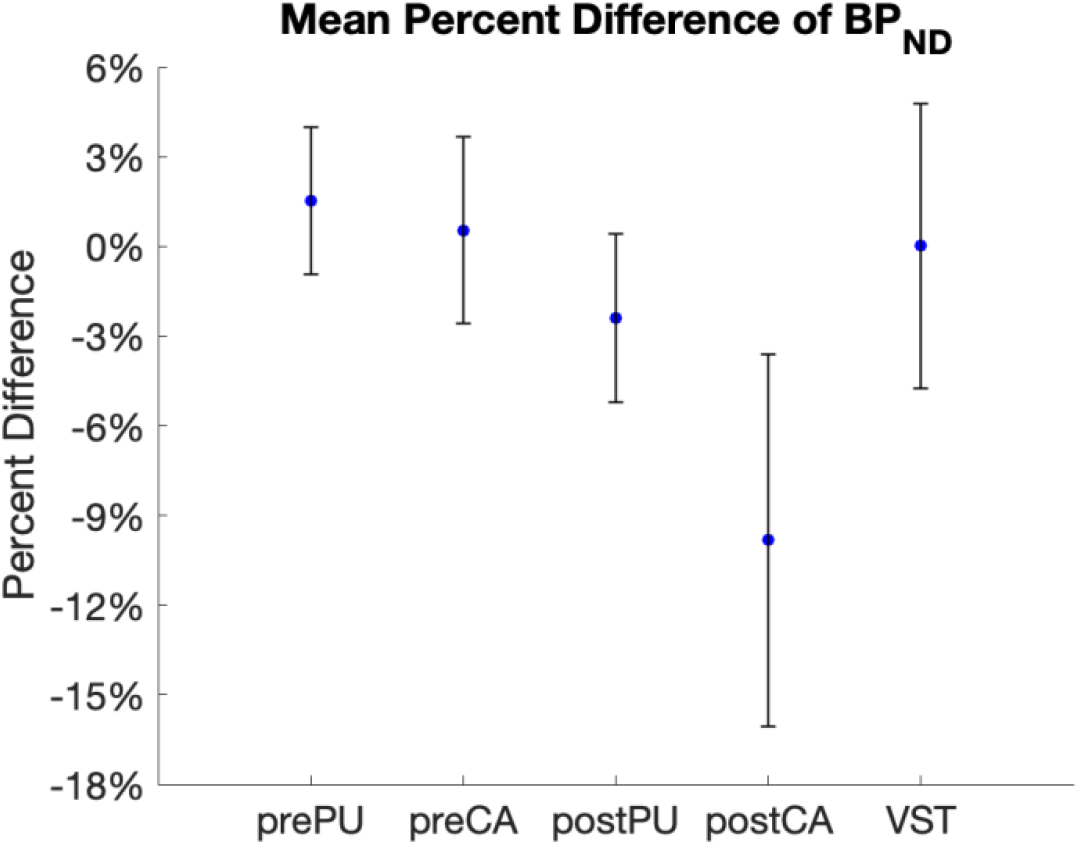
Mean Percent Differences (standard deviation) between BP_ND_ of the 5 subregions calculated using manual ROIs and MTL-generated ROI across 19 independent test subjects.

**Table 2:**
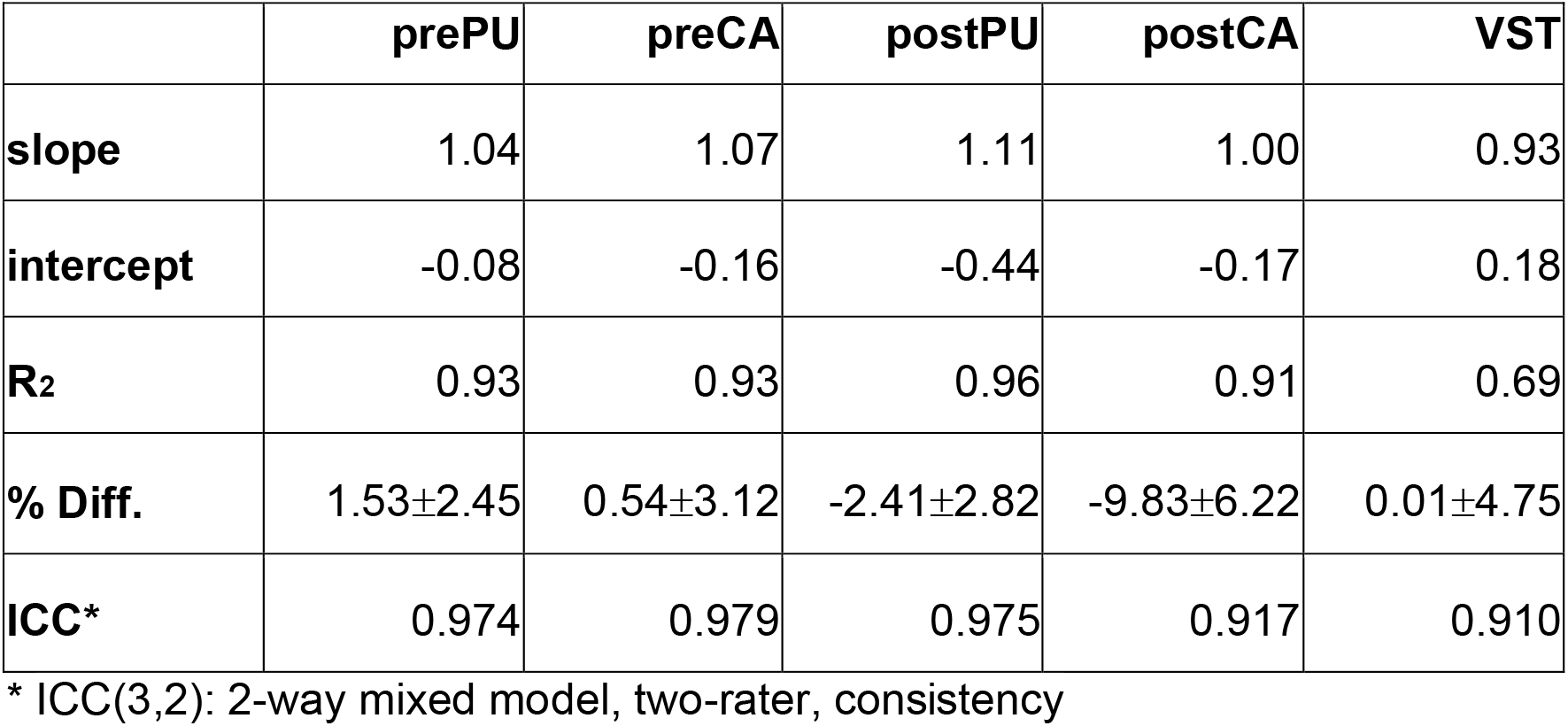
Regression coefficients from correlative analysis between BP_ND_ calculated from manually drawn ROIs and MTL-generated ROIs. R_2_ from regression analysis, percent difference with standard deviations, and ICC comparing BP_ND_ from CNN generated and manual segmentations are also shown.

|  | **prePU** | **preCA** | **postPU** | **postCA** | **VST** |
| --- | --- | --- | --- | --- | --- |
| **slope** | 1.04 | 1.07 | 1.11 | 1.00 | 0.93 |
| **intercept** | -0.08 | -0.16 | -0.44 | -0.17 | 0.18 |
| **$R_2$** | 0.93 | 0.93 | 0.96 | 0.91 | 0.69 |
| **% Diff.** | 1.53±2.45 | 0.54±3.12 | -2.41±2.82 | -9.83±6.22 | 0.01±4.75 |
| **ICC\*** | 0.974 | 0.979 | 0.975 | 0.917 | 0.910 |

\* ICC(3,2): 2-way mixed model, two-rater, consistency

fMRI results were similar, as shown in Table 3 and Figure 5, which show mean Pearson correlation coefficients between manual and MTL-generated segmentations for fMRI timeseries and whole-brain connectivity analysis. fMRI timeseries data prior to RSFC processing were correlated at 0.95 or above for preCA and both putamen subregions, with somewhat lower values of 0.87 and 0.85 for postCA and VST, respectively (RSFC processed timeseries data are not shown, but correlations differed by no more than 0.01 for all subregions). Whole-brain connectivity values were similarly highly correlated, with r values of 0.96 or higher for preCA and both putamen subregions, and of 0.91 and 0.88 for postCA and VST, respectively.

**Figure 5:**
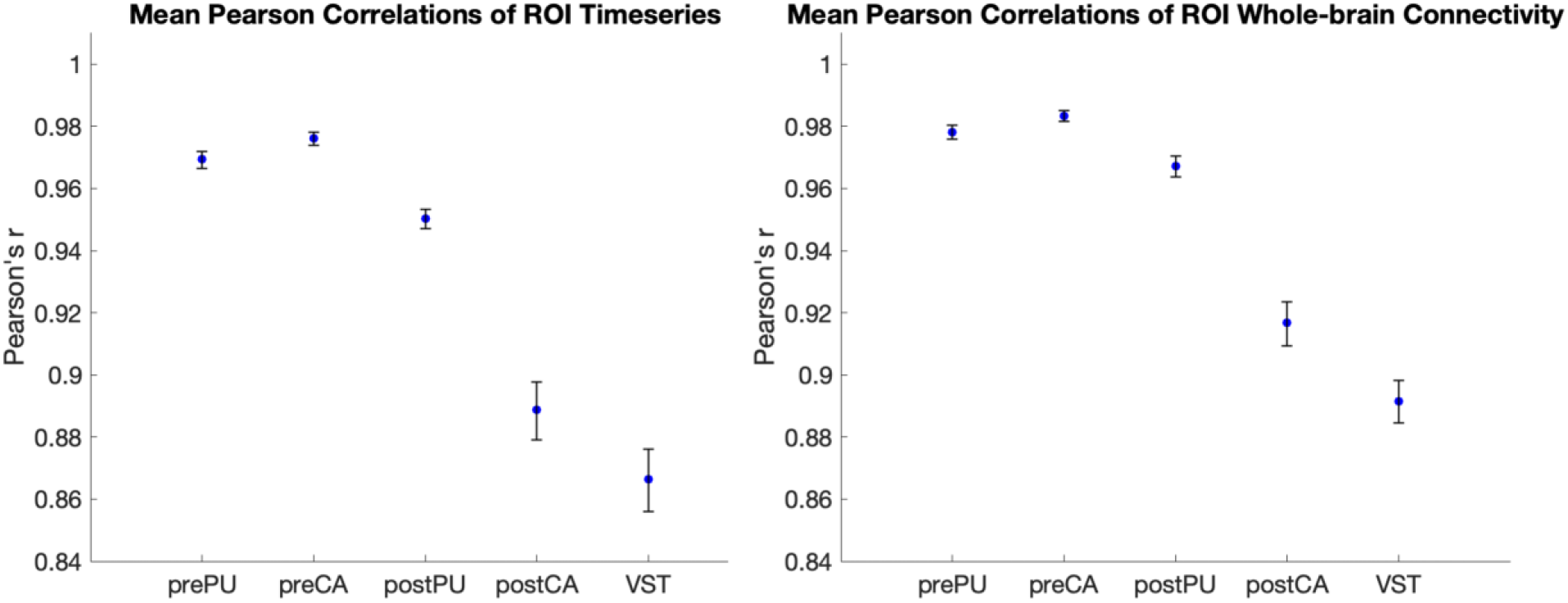
Mean Pearson correlation coefficients (Pearson’s r) with error bars depicting 95% confidence intervals between manual ROIs and MTL-generated ROIs for the all sub-regions in (left) fMRI timeseries and (right) whole-brain connectivity analysis.

**Table 3:**
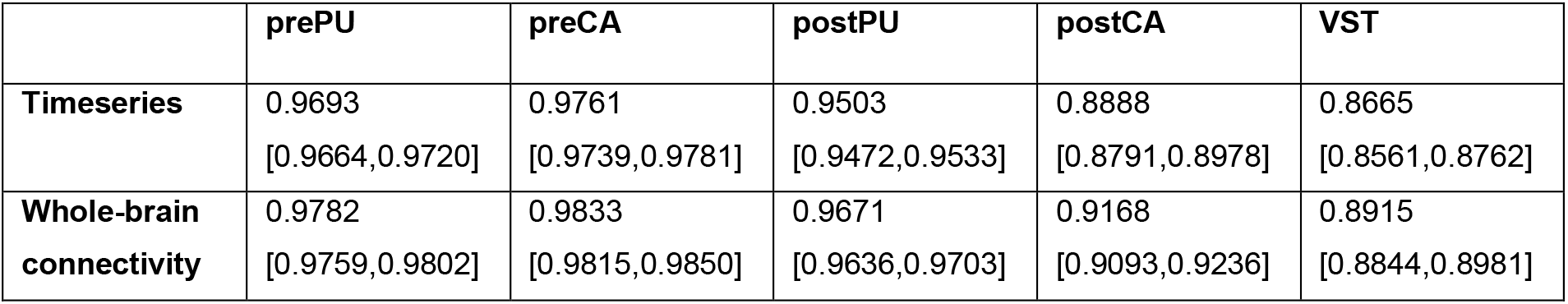
Mean Pearson correlation coefficients (Pearson’s r) with 95% confidence intervals between manual ROIs and MTL-generated ROIs for the all sub-regions in fMRI timeseries and whole-brain connectivity analysis

|  | prePU | preCA | postPU | postCA | VST |
| --- | --- | --- | --- | --- | --- |
| **Timeseries** | 0.9693<br>[0.9664,0.9720] | 0.9761<br>[0.9739,0.9781] | 0.9503<br>[0.9472,0.9533] | 0.8888<br>[0.8791,0.8978] | 0.8665<br>[0.8561,0.8762] |
| **Whole-brain connectivity** | 0.9782<br>[0.9759,0.9802] | 0.9833<br>[0.9815,0.9850] | 0.9671<br>[0.9636,0.9703] | 0.9168<br>[0.9093,0.9236] | 0.8915<br>[0.8844,0.8981] |

## 4. Discussion

In this study, we developed a deep MTL framework for striatal subregion segmentation and assessed the network outputs with PET and fMRI-derived quantitative outcome measures. We show here that this method provides extremely fast and reliable segmentations for the purpose of PET quantification, fMRI timeseries extraction, and whole-brain RSFC correlations, of the subregions of the striatum that avoid the labor-intensive task of hand drawing ROIs. This network architecture was designed to optimize the segmentation of multiple striatal subregions trained and validated on a total of 68 3D T1-weighted MR images and their manual segmentations. The network was further evaluated on 19 test datasets for PET and 15 test datasets for fMRI.

Visually, the MTL-generated ROIs look extremely similar to manual segmentations, despite smoothing of edges in the CNN output. DSC scores obtained from the comparison of manual and MTL-generated segmentations were > 0.8 in 3 of the 5 ROIs and > 0.7 across all subregions, with the lowest DSC score associated with the VST. The lower VST score is likely due to its small size and irregular shape. Nevertheless, comparative PET quantitative analysis between manual and MTL-generated segmentations showed that, although slightly smoothed, our MTL-generated segmentations agree well with the manual segmentation results. Likewise, we observe a downward bias in the postCA, a region that is long, narrow, and surrounded by background, which may make it difficult for the network to discern and segment this region. Nevertheless, R_2_ of BP_ND_ between methods was 0.91, and whole-brain RSFC of this region was correlated at 0.91 between manual and CNN ROIs, indicating that PET quantification and RSFC analysis using our network can explain most of the variance in the manually delineated data.

Growing interest in segmenting the striatum into its functional subdivisions led to a previous investigation comparing manual and automatic striatal subregion segmentation for interrater, and intrarater, reliability on measuring BP_ND_ from [11C]raclopride PET_32_. This study divided the striatum into 3 subdivisions: limbic striatum, consisting of VST; associative striatum, consisting of preCA and prePU; and sensorimotor striatum, consisting of postCA and postPU. This study not only showed large differences between two manual raters, but also showed conventional automated methods such as FreeSurfer to have high test-retest variability (TRV) of up to −15.6% for VST BP_ND_ alone. Interrater reliability between BP_ND_ measured from FreeSurfer limbic striatum segmentations and manual segmentation showed Pearson’s r of 0.81. Across associative and sensorimotor striatum, automated methods perform rather well with Pearson’s r > 0.88 and 0.95_32_, respectively. We used post-hoc analysis to group our five subregions into limbic, associative and sensorimotor functional groups as mentioned in this publication. Our results showed Pearson’s r to be 0.832 in limbic striatum, 0.96 in both substructures of the associative striatum and 0.95 and 0.98 in the substructures of the sensorimotor striatum. This suggests that our MTL-generated segmentations can be used to generate reliable measurements compared to manually based segmentations.

Fast and reliably obtained striatal subregion segmentations are extremely beneficial in research settings, facilitating accurate and reproducible quantification of neuroreceptors, indices of neurotransmission capacity and striatal-cortical connectivity_1,24,25_; all of which are of strong interest in studies of patients with schizophrenia, as well as many other neuropsychiatric and neurological disorders. Overall, fast computational speed for striatal segmentations increases feasibility of large brain imaging cohorts for large-scale structural and functional brain studies. The accuracy that our CNN-based segmentation has achieved allows this method to be a very advantageous approach for imaging studies.

An issue that all deep learning models face is the number of training/validation samples used to optimize the network. Our network used 68 subjects to train/validate the network, which is larger than the 13 datasets for deep learning striatal segmentation used in prior work_9_. Moreover, our network was able to generate segmentations for subregions of the striatum, which has not been attempted previously, to our knowledge. Using an MTL approach allows our network to share similar parameters across each ROI from the same T1-weighted input to optimize segmentation, thereby creating a more robust network for segmenting individual ROIs, despite the number of training samples used. Nevertheless, our approach could always benefit from more training datasets to improve performance and avoid the smoothing that the network has shown for the ROIs, especially VST. Other non-deep learning methods did not require manual segmentations and were able to segment regions such as the putamen and caudate_33_, using multimodal imaging such as T1-, T2-, and Diffusion-weighted imaging (DWI) from the Human Connectome Project (HCP) dataset. This DWI acquisition from HCP had a total acquisition time of one hour, which is not standard across all MRI protocols. Our network is much more generalizable since it utilizes only T1-weighted images to segment several subregions of the striatum. Nonetheless, the network used here can be easily modified to adapt to include multiple contrasts as input channels. This MTL network is designed to optimize such adaptations by utilizing hard parameter sharing between inputs to segment multiple ROIs at once. This network can also be made to segment other structures of the brain due to the flexibility inherent in our MTL network, especially since it is already trained to segment regular and irregular shapes such as the caudate and ventral striatum. Modifications and re-training using different contrasts and datasets would be required but could easily be implemented for this MTL network.

In this paper, we have developed a fast, robust and reliable method for 3D striatal subregion segmentation using an MTL approach. This deep learning performance was comparable to manually segmented ROIs but exponentially faster. Moreover, our MTL-generated segmentations were reliable in PET and fMRI quantification. Due to its speed and accuracy in segmenting striatal subregions, this MTL network has broad applicability in neuroimaging.

## 5. Acknowledgement

The authors would like to thank Xiaoyan Xu PhD for performing the manual segmentation on some of the data sets. Previously published data used in Training Set 1 was funded by P50 MH 086404 (AAD). Unpublished data used in Training Set 2 was funded by K01 MH 107763 (JXVS). Previously published data used in Testing Set 1 was funded by R21 MH 099509 (AAD). JJW was supported by K23 MH 115291, JXVS was supported by K01 MH 107763, and CH was supported by NARSAD 24971.

